# Parallel and Divergent Evolution in *Pseudomonas aeruginosa* Under Constant and Fluctuating Predator-Mediated Selection

**DOI:** 10.64898/2026.07.22.739916

**Authors:** Nayma Romo Bechara, Marcos Garcia, Michael J. Bland, Kasie Raymann

## Abstract

Environmental predation is a major driver of bacterial evolution and may indirectly influence virulence through coincidental selection. However, how sustained versus fluctuating predator pressure shapes long-term evolutionary trajectories remains poorly understood. Here, we used experimental evolution to investigate the genetic and phenotypic responses of *Pseudomonas aeruginosa* to continuous, absent, or fluctuating exposure to the protozoan predator *Tetrahymena thermophila* over 180 days. Whole-population and isolate-level shotgun metagenomic sequencing revealed fewer mutations over time but increasing frequencies of surviving mutations, consistent with selection, extensive gene-level parallel evolution, and signatures of both positive and purifying selection. Recurrently mutated genes encompassed diverse functional pathways, reflecting both shared and treatment-specific adaptive responses. Despite this parallelism, historical contingency was evident, with starting conditions influencing subsequent evolutionary trajectories. We also observed the emergence of hypermutator lineages, which are frequently recovered from chronic lung infections, suggesting that repeatedly evolving elevated mutation rates may represent a common adaptive strategy of *P. aeruginosa* across environmental and host-associated settings. Fluctuating predation repeatedly reshaped the adaptive landscape, leading to greater temporal turnover of mutations and a higher accumulation of mutations that ultimately reached fixation than in constant environments. Phenotypic assays revealed widespread divergence in fitness, motility, biofilm formation, siderophore production, protease activity, hemolysis, and cell size, whereas virulence in an invertebrate host model varied among treatments but did not differ significantly. Together, these findings demonstrate that variation in predator-mediated selection reshapes the dynamics and genetic targets of bacterial adaptation, highlighting the roles of ecological context, historical contingency, and hypermutability in driving the evolutionary trajectories of opportunistic pathogens.

**Significance Statement:** Environmental predators are drivers of bacterial evolution, yet their effects on adaptation remain poorly understood. We used experimental evolution to show that constant and fluctuating protozoan predation produce evolutionary trajectories in Pseudomonas aeruginosa, altering tempo, predictability, and targets of adaptation. Adaptation to predator-present or predator-absent environments shaped evolutionary trajectories, demonstrating importance of historical contingency. Fluctuating predation promoted turnover of mutations as populations adapted to selective pressures. We also observed repeated emergence of hypermutator lineages, a hallmark of chronic infections, suggesting that elevated mutation rates represent a favored adaptive strategy across environmental and host-associated settings. These findings provide insight into the environmental origins of genetic changes commonly associated with opportunistic pathogens, while showing that these changes do not necessarily increase virulence.

## Introduction

*Pseudomonas aeruginosa* is a model organism in evolutionary microbiology because of its remarkable ecological versatility, genetic plasticity, and clinical importance (1–3). It inhabits diverse environmental niches, including soils, aquatic systems, plant surfaces, and host-associated environments, while also causing persistent infections, particularly in individuals with cystic fibrosis and chronic obstructive pulmonary disease (4–6). Adaptation to these contrasting environments has made *P. aeruginosa* an important system for understanding how ecological selection shapes bacterial evolution. Notably, hypermutator lineages frequently emerge during chronic infections (7–10) accelerating adaptation to host-associated selective pressures, although whether elevated mutation rates also represent a common strategy during environmental evolution remains unknown.

Predation is a pervasive ecological force shaping microbial communities, with bacterivorous protozoa imposing strong selection for traits such as altered motility, biofilm formation, toxin production, and surface modification that enhance resistance to grazing (11–13). Because many of these adaptations also contribute to host colonization and persistence, environmental predation has been proposed as an important driver of coincidental virulence evolution (12,14). However, evolution under protist predation has been examined in *P. aeruginosa* in only two short-term experimental evolution studies: our previous 60-day evolution experiment (15) and that of Leong et al. (16). Together, these studies demonstrated that predator-mediated selection can alter adaptive trajectories and virulence-associated phenotypes but were limited to constant predator environments over relatively short evolutionary timescales. In natural ecosystems, predator pressure is rarely constant. Instead, bacterial populations experience alternating periods of grazing and predator absence that repeatedly alter selective pressures and may redirect adaptive trajectories, maintain genetic diversity, and strengthen the effects of historical contingency (17,18). Consequently, it remains unknown whether the evolutionary patterns observed during short-term, constant predator exposure persist during prolonged evolution or how fluctuating predator pressure influences long-term adaptation.

To address these questions, we evolved *P. aeruginosa* populations for 180 days under continuous predator exposure, predator-free conditions, or alternating predator and predator-free environments using the ciliate *Tetrahymena thermophila* as a predator. Fluctuating populations experienced an initial 60-day period in either predator or predator-free conditions before alternating every two weeks, allowing us to evaluate the effects of pulsed predation pressure and historical contingency on subsequent adaptation. Whole-population and isolate-level shotgun metagenomic sequencing revealed extensive parallel evolution, signatures of positive and purifying selection, and treatment-specific adaptive responses across diverse functional pathways. We further identified recurrent evolution of hypermutator lineages under environmental selection, suggesting that elevated mutation rates represent a broadly deployed adaptive strategy in *P. aeruginosa* rather than one restricted to chronic infections. Complementary phenotypic analyses revealed widespread divergence in fitness, motility, biofilm formation, siderophore production, protease activity, hemolysis, and cell size, whereas virulence in an invertebrate host model remained largely unchanged. Together, these integrated genomic and phenotypic analyses demonstrate how ecological context, historical contingency, and recurrent hypermutability shape the evolutionary trajectories of *P. aeruginosa* under constant and fluctuating predator-mediated selection.

## Results

### Genotypic Analysis of Evolved Populations

Replicate populations of *P. aeruginosa* NRB were evolved for 180 days under two stable conditions: predator-free media (ME, media-evolved; n = 6) and media containing the protozoan predator *T. thermophila* (PE, predator-evolved; n = 6). The initial 60 days of evolution were reported previously (15). Two fluctuating treatments were established by transferring populations that had evolved for 60 days in either predator-free media or media containing *T. thermophila* to alternating environments at two-week intervals for the remaining 120 days: PM (predator-to-media; n = 6) and MP (media-to-predator; n = 6). All populations were sampled at days 60, 120, and 180 and subjected to whole-genome shotgun metagenomic sequencing. Each evolved population was compared to the ancestral genome. One ME and one MP population were removed from downstream analyses due to contamination detected at Day 60 (15). Three independently evolved hypermutator lineages (ME6, PE5, and PM5) emerged during the experiment (**Figure 1A–B, Dataset S1**), driving the marked variation in mutational burden across populations (7–696 mutations; minimum frequency cutoff = 0.05; mean = 79). The number of mutations per population was not correlated with average genome coverage (**Figure S1, Table S1**), indicating that sequencing depth did not influence mutation detection.

**Figure 1.**
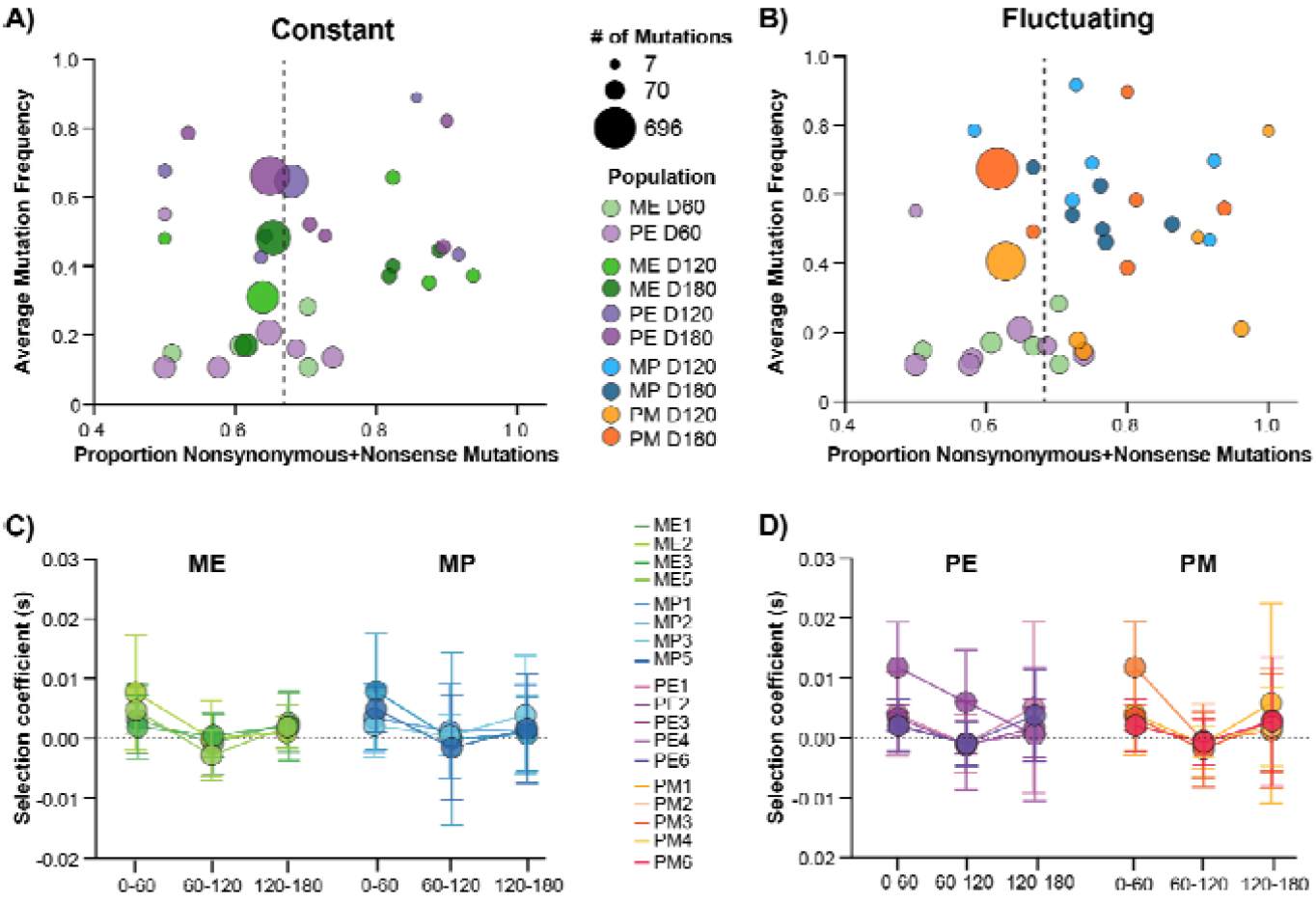
Mutation frequency, mutation composition, and selection coefficients across experimentally evolved populations. **(A)** Populations evolved under constant media (ME) or predator exposure (PE). **(B)** Populations evolved under fluctuating environments, beginning in media (MP) or predator (PM) for 60 days before alternating between predator and predator-free conditions every two weeks. Scatter plots show the average mutation frequency (y-axis) versus the proportion of nonsynonymous and nonsense mutations (x-axis) for each population at Days 60, 120, and 180. Circle size is proportional to the total number of mutations detected per population. The vertical dashed line indicates the expected proportion of nonsynonymous and nonsense mutations under neutrality. **(C–D)** Mean selection coefficient(s) of all detected mutations across three evolutionary intervals (Days 0– 60, 60–120, and 120–180) for each replicate population. Error bars represent the standard deviation of selection coefficients within each population. The horizontal dashed line denotes neutrality (s = 0), with positive values indicating increasing allele frequencies and negative values indicating decreasing allele frequencies. Importantly, s = 0 does not necessarily imply neutrality or the absence of selection, as mutations that have already reached fixation (or are otherwise maintained at a constant frequency) will also have s = 0 despite having been subject to selection. Colors identify individual replicate populations.

All evolved populations contained nonsynonymous mutations, with most also harboring synonymous, intergenic, and nonsense mutations, while a subset contained mutations in noncoding (RNA) regions (**Table S2, Dataset S1**). Across both stable and fluctuating environments, average mutation frequencies increased over time, with Day 120 and Day 180 populations exhibiting higher frequencies than Day 60 populations (**Figure 1A–B**). Comparison with neutral expectations generated from 10,000 simulated mutations revealed that most Day 60 populations contained fewer nonsynonymous mutations than expected, whereas nearly all Day 120 and Day 180 populations exhibited an excess of nonsynonymous mutations, consistent with strong positive selection (**Figure 1A–B, Table S2**). In contrast, the three hypermutator populations (ME6, PE5, and PM5) generally contained fewer nonsynonymous mutations than expected, suggesting that purifying selection removed many protein-altering mutations despite their elevated mutation rates. Unless otherwise stated, all subsequent analyses excluded the hypermutator populations.

Selection coefficients, calculated for each detected mutation based on variation in allele frequency between sampling intervals, varied among treatments and over time (**Figure 1C-D, Dataset S2**). Across all treatments, selection coefficients were generally highest during the first sampling interval (Days 0–60), declined between Days 60 and 120, and increased again during the final interval (Days 120–180). Although this overall trend was observed across all treatments, populations that experienced predator exposure (PE, MP, and PM) exhibited substantially greater temporal variation in selection coefficients during the later sampling intervals than populations maintained in the no-predator control (ME). In particular, PE, MP, and PM populations contained more mutations exhibiting both strongly positive and strongly negative selection coefficients between sampling intervals whereas ME populations generally showed less variation in selection coefficients and fewer changes in allele frequencies over time (Figure 1C-D).Because selection coefficients were estimated across 60-day intervals, these values are conservative and likely underestimate the maximum strength of selection experienced by mutations that rapidly swept to fixation within an interval.

To identify the mutations underlying these temporal patterns, we examined alleles exhibiting the strongest positive and negative selection coefficients across all detected mutations (**Figure 2**). Recurrently selected mutations were concentrated in a small number of genes, including *phoQ, marR, mgtE, fleQ, dipA*, and *wbpM*, demonstrating extensive parallel evolution across replicate populations. Mutations with selection coefficients of zero generally represented alleles that had already reached fixation and whose frequencies remained unchanged across subsequent sampling intervals. Because selection coefficients are estimated from changes in allele frequency over time, once a mutation is fixed (or otherwise remains at a constant frequency), there is no detectable frequency change from which to estimate ongoing selection. Despite the fixation of many early adaptive mutations, new mutations continued to arise and sweep through populations over the course of the experiment, particularly in populations exposed to predators.

**Figure 2.**
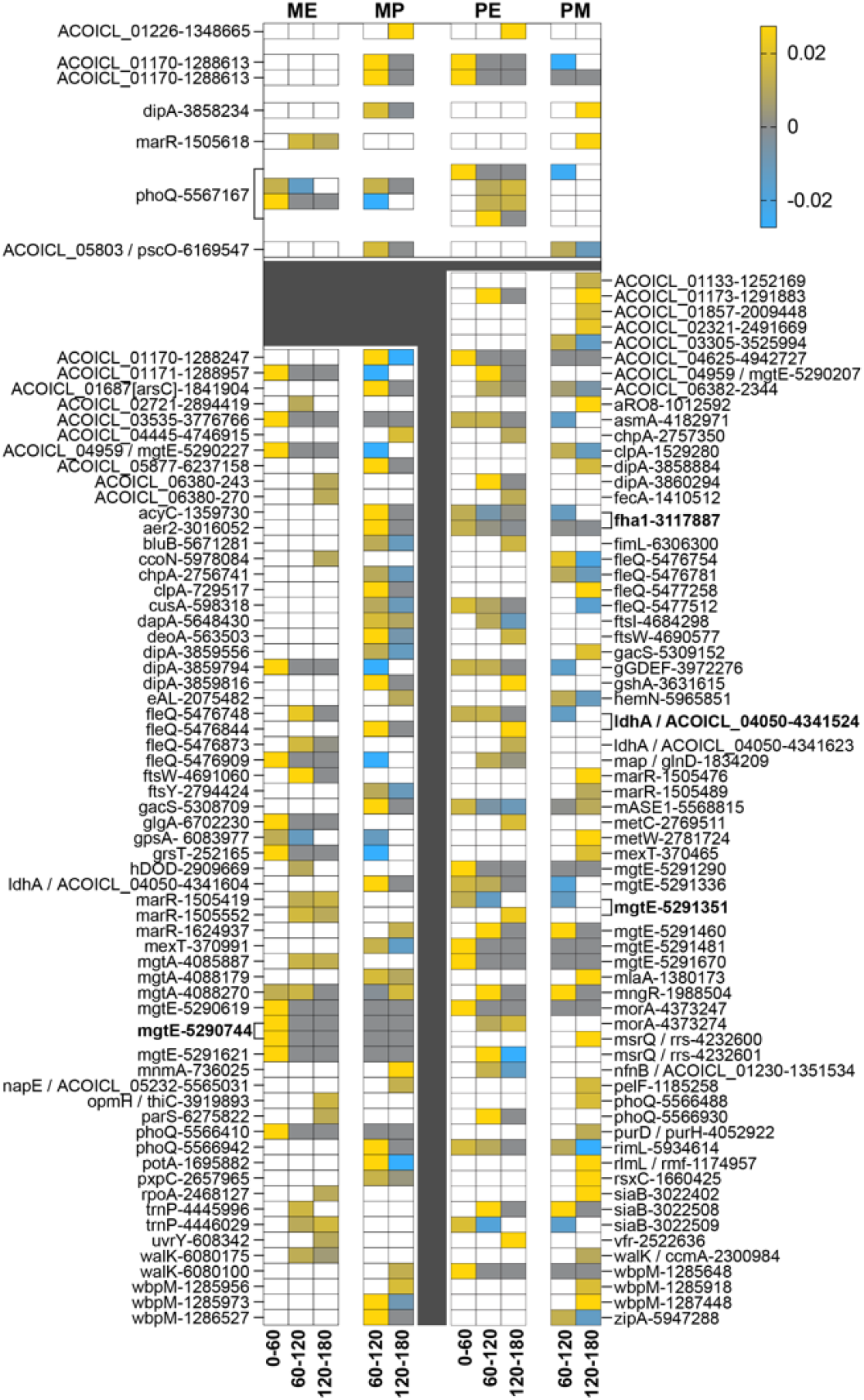
Heatmap of selection coefficients (s) for alleles exhibiting the strongest positive and negative selection during experimental evolution. Columns represent the four treatments (ME, MP, PE, and PM), with selection coefficients calculated across consecutive evolutionary intervals: Day 0–60, Day 60–120, and Day 120–180. Yellow indicates positive selection, blue indicates negative selection, gray indicates a selection coefficient of approximately zero (s ≈ 0), meaning the mutation did not change in frequency over the measured interval, and white indicates that the mutation was absent from the population at the corresponding time point. Mutation labels indicate the affected gene (or intergenic region) followed by the genomic position of the mutation. The **upper panel** (above the dark gray divider) highlights mutations occurring at the same genomic position across selection regimes (ME-PE, MP-PE, PM-ME, PM-MP). The **lower panel** contains strongly selected mutations restricted to the ME and/or MP populations or the PE and/or PM populations and therefore not shared across both treatment groupings. Bolded labels enclosed by brackets identify genomic positions that were independently mutated in two or more populations, either within the highlighted upper panel or among the remaining mutations in the lower panel.

Analysis of mutation frequency distributions revealed a strongly bimodal pattern, with most mutations occurring either at very low frequencies (0.05– 0.1) or approaching fixation (>0.95) (**Figure 3A**). To determine whether mutations reaching high frequency represented a distinct class of variants, we compared the distribution of nonsynonymous, nonsense, synonymous, intergenic, and RNA mutations among all detected variants and among mutations reaching ≥0.5 frequency in at least one population against neutral expectations. Whereas the distribution of all detected mutations (≥0.05 frequency) did not differ substantially from neutral expectations, high-frequency mutations were strongly enriched for nonsynonymous and nonsense variants, which comprised 79.3% of these mutations compared to ∼67% expected under neutrality (**Figure 3B**). Together with the selection coefficient analyses, these findings indicate that mutations reaching high frequency were the mutations most strongly favored by positive selection and, consequently, the variants most likely to contribute to adaptive phenotypic change. Therefore, all subsequent analyses were restricted to high-frequency mutations (≥0.5), yielding a dataset of 134 mutations (**Figure S2, Dataset S3**). Most high-frequency mutations were single nucleotide polymorphisms, with the remainder comprising 19 deletions, eight duplications, and one insertion. Although most deletions involved only 1–3 bp, larger deletions ranging from 12 bp to 42.9 kb were also observed. Many affected genes that also accumulated high-frequency SNPs, including *ftsY, siaB, mgtE*, and *fleQ*. In MP2, an exceptionally large 42.9-kb deletion encompassing 62 phage-associated genes reached fixation by Day 120 and persisted through Day 180, indicating stable loss of the prophage region **(Dataset S3)**.

**Figure 3.**
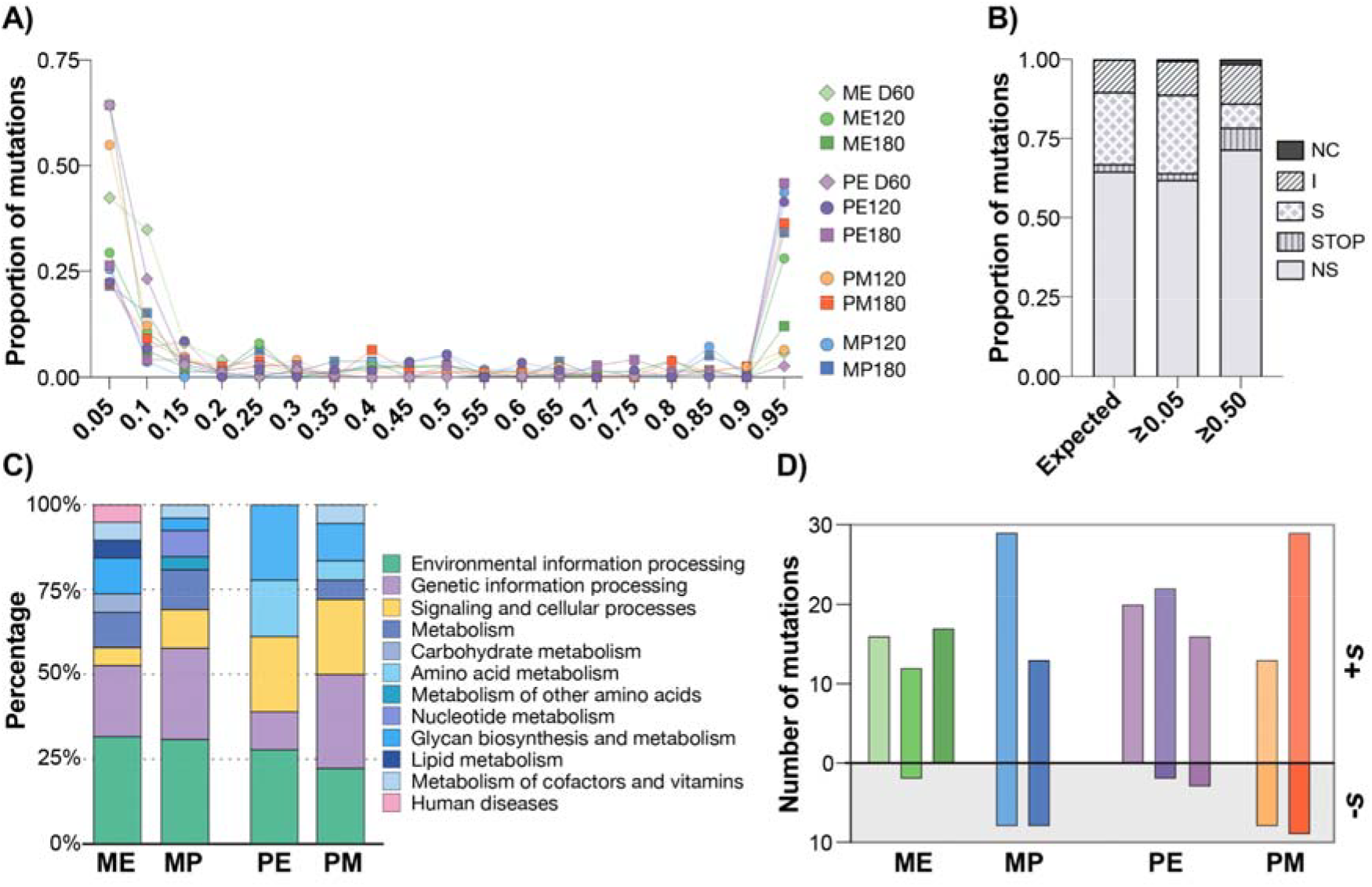
Mutation frequency distribution and proportion types of mutations. **(A)** Number of mutations detected at different frequencies between 0.5-1 across all evolved populations. Each bar represents the lower bound of each 0.05 interval (e.g., 0.05 = 0.05–0.09; 0.95 = 0.95– 1.0). **(B)** Proportions of nonsynonymous, nonsense, synonymous, intergenic, and RNA mutations expected under neutral evolution compared with those observed across all detected mutations (≥0.05 frequency) and among high-frequency mutations (≥0.5 frequency), excluding hypermutator populations (−H) from the latter analysis. **(C)** KEGG functional classification of high-frequency mutations (≥50%) occurring in coding regions. **(D)** Number of high frequency mutations across the three evolutionary intervals (Days 0–60, 60– 120, and 120–180) with positive (+s) or negative (-s) selection coefficients.

Among the 134 high-frequency mutations, some were unique to individual evolutionary treatments whereas others arose repeatedly across independent populations, demonstrating extensive parallel evolution (**Figure S2, Dataset S3**). We identified 10 independent site-specific parallel mutation events after excluding repeated observations within individual populations. These events were unevenly distributed across treatments, with the greatest sharing occurring between PE and PM populations, whereas no site-specific mutations were shared across all four treatments. At the gene level, 75 genes exhibited parallel evolution, including 28 mutated across all treatments. Frequently targeted genes included *clpA, marR, mgtA, gacS, wbpM, siaB, ldhA, mexT*, and *morA*, whereas repeated mutations in *trnP* and *mrsQ* were restricted to ME and PE populations, respectively (**Dataset S3**).

KEGG annotation of coding-region mutations revealed treatment-specific functional biases among adaptive mutations (**Figure 3C**). Approximately 30% of high-frequency mutations in ME, PE, and MP populations affected environmental information processing compared with 22% in PM populations. Predator-exposed populations (PE, PM, and MP) were enriched for mutations affecting signaling and cellular processes, whereas ME populations contained relatively more mutations associated with metabolism. Distinct metabolic signatures were also evident: carbohydrate and lipid metabolism mutations were unique to ME, amino acid metabolism mutations occurred only in PE and PM, and mutations affecting cofactor and vitamin metabolism were restricted to fluctuating populations (MP and PM), highlighting divergent adaptive trajectories among evolutionary regimes (**Figure 3C**).

To further examine the temporal dynamics of adaptive evolution, we quantified the number of high-frequency mutations exhibiting positive or negative selection coefficients during each evolutionary interval (**Figure 3D**). Across all treatments, mutations with positive selection coefficients greatly outnumbered those with negative coefficients, indicating that most high-frequency mutations were under positive selection. However, the temporal distribution of mutations with positive estimated selection coefficients differed among treatments. The fluctuating predator treatments (MP and PM) exhibited distinct temporal spikes in the number of positively selected mutations, with MP showing the strongest signal during the first interval (Days 60–120) and PM during the final interval (Days 120–180). In contrast, the constant treatments (ME and PE) maintained relatively steady numbers of positively selected mutations across both sampling intervals. Overall, relatively few high-frequency mutations exhibited negative selection coefficients. When negative selection was observed, it occurred almost exclusively in the fluctuating predator treatments (MP and PM), whereas the constant treatments (ME and PE) showed little to no evidence of negatively selected mutations. These results indicate that fluctuating environments more frequently changed the direction of selection on existing variants, leading to episodic turnover among high-frequency alleles.

The temporal dynamics of high-frequency mutations further distinguished the evolutionary treatments (**Figure S2**). Several mutations that arose by Day 60 persisted throughout the experiment under both constant and fluctuating conditions, including mutations in *mgtE, phoQ, wbpM, morA*, and a NAD-dependent dehydratase. However, additional high-frequency mutations continued to arise, particularly in fluctuating populations, where 13 and 11 new mutations arose in MP and PM populations, respectively, compared with only four in PE populations. In addition, two early high-frequency mutations were lost exclusively under fluctuating conditions, and five mutations exhibited transient dynamics, appearing at Day 120 but disappearing by Day 180. Together, these temporal patterns complement the selection coefficient analyses **(Figures 2 and 3D)** by showing that fluctuating environments generated episodic bursts of positive selection and greater turnover among high-frequency mutations, whereas constant environments maintained a more stable set of high-frequency variants through time.

### Genotypic and Phenotypic Analysis of Evolved Isolates

To determine how genomic evolution translated into phenotypic divergence, single isolates were randomly selected from 18 populations at Day 180 for genotypic and phenotypic characterization (see Methods for selection criteria). Across the 18 isolates included in the analysis, we identified 119 distinct mutations, with individual isolates carrying 4–13 mutations (mean = 8.5 mutations per isolate; **Figure S3, Dataset S4**). Nearly all mutations detected in the isolates were fixed within their corresponding populations, indicating that they were representative of the dominant population genotype. We quantified swimming and swarming motility, biofilm formation, protease activity, hemolysis, siderophore production, cell size, growth, virulence in an invertebrate host model, and competitive fitness.

Distinct phenotypic trajectories emerged among evolutionary treatments (**Figure 4, Figure S4**). Isolates evolving under fluctuating predator regimes (PM and MP) consistently exhibited reduced swimming motility, biofilm production, siderophore production, and cell size relative to the ancestor, indicating highly repeatable responses to fluctuating selection. In contrast, isolates evolving under constant environments (ME and PE) generally displayed increased swarming motility and siderophore production, although biofilm production varied among replicate populations. Hemolytic activity showed one of the most consistent treatment-specific responses, with all PM isolates exhibiting significantly greater activity than the ancestor. Together, these results demonstrate that fluctuating and constant predator regimes promoted distinct trajectories of phenotypic adaptation despite extensive parallel evolution at the genomic level.

**Figure 4.**
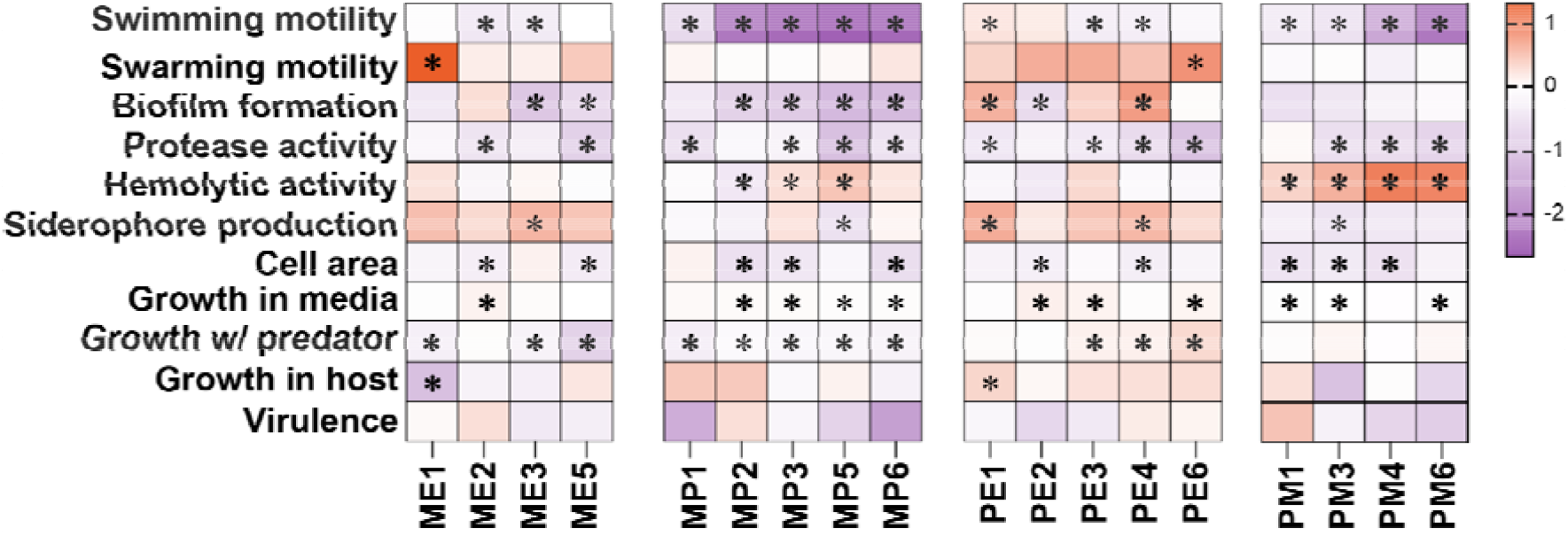
Log_2_ fold change in phenotypes relative to the ancestral strain. Asterisks indicate statistically significant differences based on one-way ANOVA with Holm–Šídák’s multiple comparisons test, using all raw values (see Figure S2). Bold asterisks denote P < 0.001, and non-bold asterisks denote P < 0.05.

Growth responses likewise depended on evolutionary history (**Figure 4, Figure S4)**. Growth in predator-free media increased in nearly all evolved isolates, consistent with adaptation to laboratory conditions. However, growth in the presence of predators diverged among treatments: it decreased in ME and MP isolates, increased in PE isolates, and remained largely unchanged in PM populations, indicating that adaptation to fluctuating predator regimes did not simply mirror adaptation to continuous predator exposure.

Despite widespread divergence in virulence-associated phenotypes, these changes did not translate into increased pathogenicity. Virulence in an invertebrate host model (*Apis mellifera*; honey bee) did not differ significantly from that of the ancestral strain across any treatment, nor did evolved isolates exhibit altered growth within the host. (**Figure 4, Figure S4**). Thus, extensive genomic and phenotypic diversification under predator-mediated selection occurred without detectable changes in host virulence.

Competition assays further demonstrated that fitness advantages were strongly environment dependent (**Figure 5**). Under laboratory conditions, competitive dominance varied among evolved isolates and depended on the environment in which they had evolved. In media alone, PE2 dominated among isolates from constant environments, whereas PM6 was the dominant competitor among isolates from fluctuating environments. In the presence of predators, PE isolates generally outcompeted ME isolates, while MP isolates collectively comprised more than 80% of fluctuating communities, with MP5 exhibiting the greatest competitive success. In contrast, the ancestral strain completely displaced all evolved isolates in the honey bee model; none remained detectable after 24 h, indicating that adaptive changes conferring competitive advantages *in vitro* did not enhance fitness within the host (**Figure 5**).

**Figure 5.**
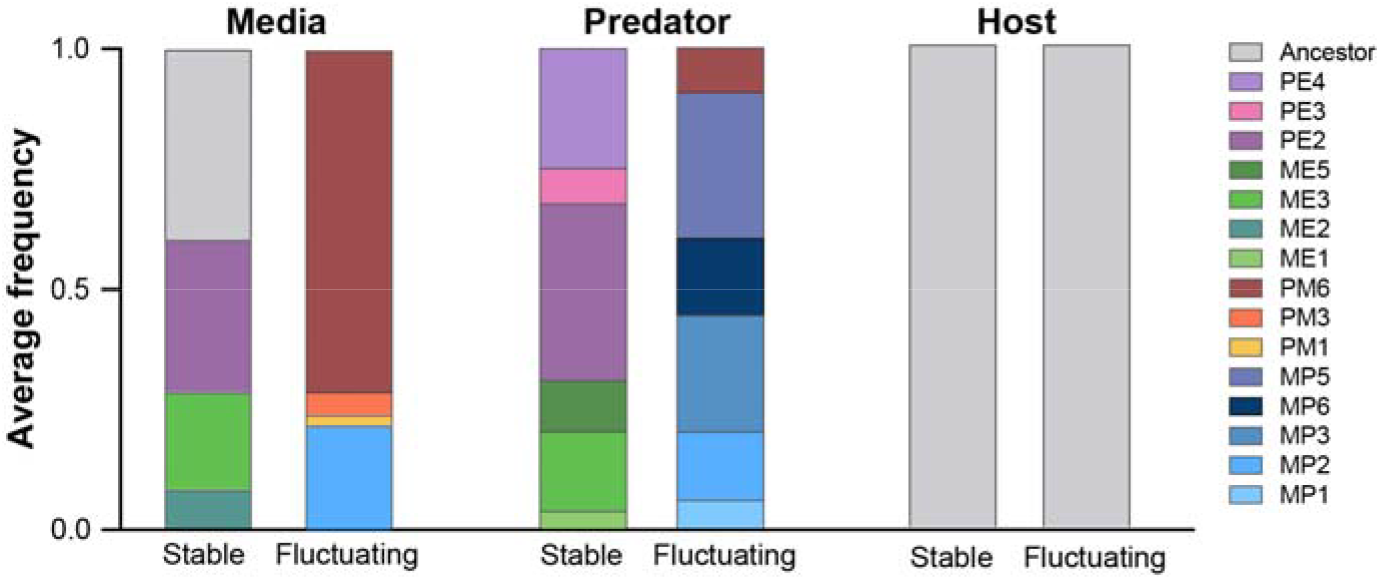
Competitive performance of evolved isolates across environments and evolutionary histories. Bars show the average frequency of each evolved isolate (except hypermutators) and the ancestral strain after 24 h of competition in media, in co-culture with *T. thermophila* (predator), or during growth in germ-free honey bees (host). Competitions were performed *separately* for isolates evolved under stable and fluctuating regimes; isolates from stable and fluctuating treatments were not pooled or competed against one another. For each competition, D180 isolates from the two treatments within each evolutionary regime were pooled in equal proportions with the ancestral strain at the start of the experiment. Values represent averages across three biological replicates, with ∼300 colonies sampled per replicate. The presence and relative frequency of each isolate after 24 h were determined by whole-genome sequencing (WGS).

### Phenotypic Analysis of Hypermutator Isolates

Whole-genome sequencing of one isolate from each independently evolved hypermutator population at D180 (ME6, PE5, and PM5) identified 202–466 mutations per isolate. Comparison with the corresponding population genomic data revealed that these isolates contained a large proportion of the mutations detected within their source populations (202 of 413 mutations in ME6, 466 of 570 in PE5, and 314 of 696 in PM5; **Dataset S5**). All three isolates contained nonsynonymous mutations in *mutS*, consistent with defective mismatch repair and the hypermutator phenotype. Although hypermutator lineages continue to accumulate mutations during routine culturing, resulting in ongoing genetic diversification, given the repeated emergence of hypermutators in our experiment and their frequent occurrence during chronic *P. aeruginosa* infections, we characterized their phenotypes.

The three hypermutator lineages exhibited extensive but distinct phenotypic divergence relative to the ancestral strain (**Figure 6, Figure S5**). Despite their elevated mutation rates and the expectation of continued mutation accumulation during routine culturing, phenotypic variation among biological replicates was generally modest **(Figure S5)**, with most phenotypes remaining reproducible despite ongoing genetic diversification. Swarming motility was significantly increased in ME6, whereas swimming motility was significantly reduced in PM5. Hemolytic activity was significantly decreased in PE5 but increased in PM5, while biofilm production increased only in ME6. Protease activity was significantly reduced in both PE5 and PM5, and siderophore production was significantly increased in ME6 and PE5 but not PM5. Growth also differed substantially among hypermutator lineages (**Figure 6A, Figure S5**). In predator-free media, all three lineages exhibited significantly greater growth than the ancestral strain. In the presence of predators, ME6 displayed significantly reduced growth, whereas PE5 and PM5 exhibited significantly increased growth relative to the ancestor. Within the invertebrate host, only PM5 showed significantly reduced bacterial abundance. Despite these differences in growth and multiple virulence-associated traits, none of the hypermutator lineages exhibited significantly altered virulence relative to the ancestral strain (**Figure 6A, Figure S5**).

**Figure 6.**
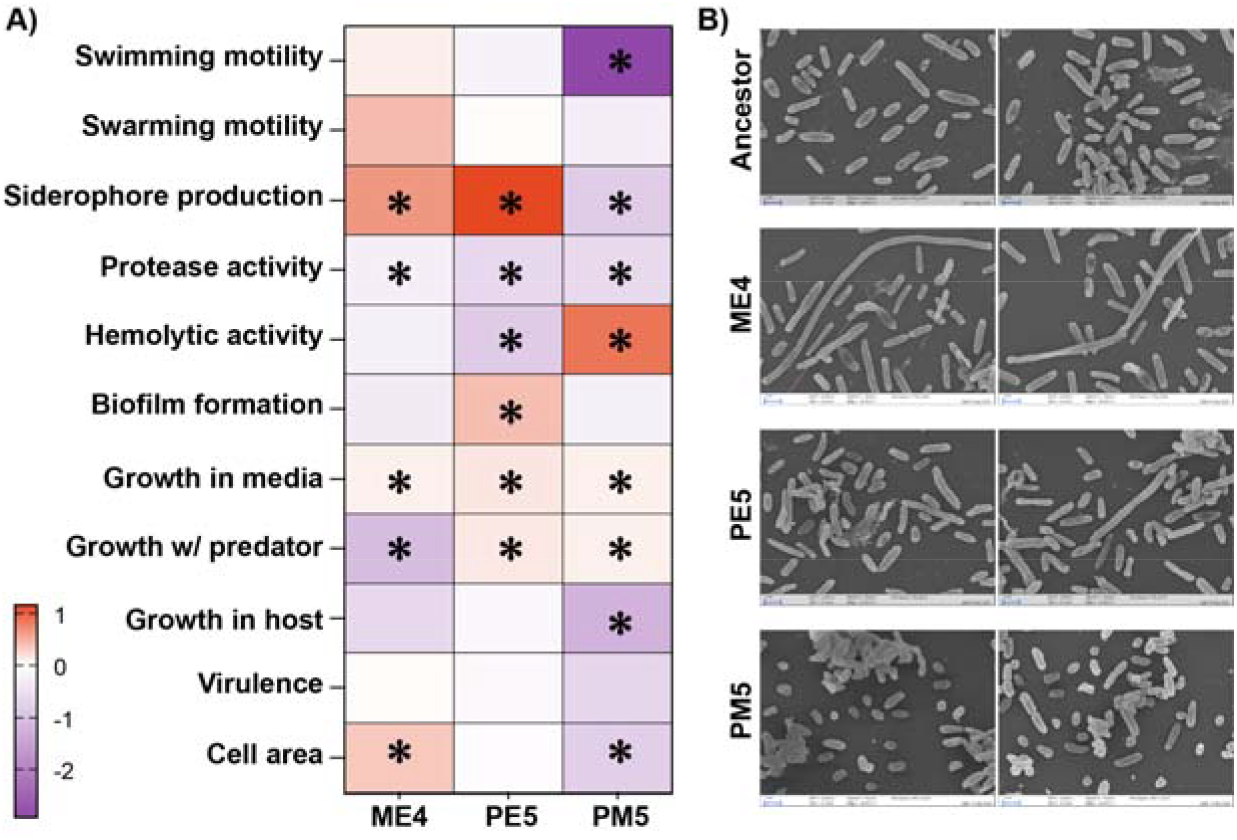
Phenotypes of hypermutator lineages. **(A)** Heatmap showing Log2Fold change of each tested phenotype for the three hypermutator lineages at D180 relative to the ancestral strain. Statistical significance relative to the ancestral strain was determined by one-way ANOVA with Holm–Šídák’s multiple comparisons test using all raw values and is indicated as *P* ≤ 0.05 (*), *P* ≤ 0.001 (**), and *P* < 0.0001 (***) (**See Figure S5**). All phenotypic assays were performed with at least three biological replicates, and all individual data points from replicate experiments are shown in **Figure S5. (B)** SEM images of hypermutator isolates (https://www.ebi.ac.uk/biostudies/bioimages/studies/S-BIAD2916).

Scanning electron microscopy revealed pronounced morphological diversification among the hypermutator populations (**Figure 6A-B**). Average cell area decreased significantly in PM5 and increased significantly in ME6 (**Figure 6A, Figure S5**). PM5 consisted predominantly of smaller cells, whereas ME6 contained many extremely elongated cells. In addition, both PE5 and ME6 exhibited substantial heterogeneity in cell size and morphology (**Figure 6B**). Whole-genome sequencing further revealed mutations affecting diverse regulatory and cellular functions in these populations (**Dataset S5**).

## Discussion

Environmental predation is a major ecological force shaping bacterial evolution, yet how temporally stable versus fluctuating predation influences long-term adaptive trajectories has remained largely unexplored. By extending our previous 60-day evolution experiment (15) to 180 days and incorporating fluctuating predator regimes, we show that predator-mediated selection influences not only the genetic targets of adaptation but also the evolutionary pathways through which adaptation proceeds. Across all treatments, populations exhibited continued parallel evolution, signatures of strong positive selection, and the repeated emergence of hypermutator lineages, indicating that adaptation remained dynamic over the course of the experiment. Similar patterns of continued adaptation have been observed in the Long-Term Evolution Experiment (LTEE), where *Escherichia coli* populations have continued to acquire beneficial mutations and increase in fitness over tens of thousands of generations despite evolving under a constant laboratory environment (19-22). Together, these studies demonstrate that prolonged bacterial adaptation need not approach a simple evolutionary endpoint. Fluctuating environments additionally promoted greater turnover in adaptive mutations and stronger effects of historical contingency, highlighting how temporal variation in selection shapes the evolutionary trajectories of adapting populations. These longer evolutionary timescales enabled us to track adaptive evolution over time, revealing evolutionary processes that were not apparent during short-term evolution. Despite extensive genomic and phenotypic diversification, however, predator-mediated selection did not produce a consistent increase in virulence in our invertebrate model system, indicating that predator-mediated selection reshapes the dynamics of adaptation without predictably increasing pathogenicity.

One of the clearest differences between evolutionary regimes was the way adaptive mutations accumulated through time. Populations maintained under constant predator or predator-free environments were characterized by the fixation and persistence of adaptive mutations, consistent with sustained directional selection in relatively stable environments. In contrast, populations experiencing alternating predator exposure exhibited substantially greater turnover in adaptive mutations, with previously successful alleles frequently being replaced by newly emerging variants. Such dynamics support theoretical predictions that fluctuating environments can maintain genetic diversity by continually changing the fitness effects of individual mutations (23-26).

The extended duration of the experiment also revealed a pronounced role for historical contingency in shaping evolutionary outcomes. Several adaptive mutations that arose during the first 60 days remained fixed throughout the experiment, demonstrating that early beneficial mutations continued to confer fitness advantages over prolonged evolutionary timescales. At the same time, populations evolving under fluctuating environments frequently lost previously favored mutations while acquiring novel adaptive variants, illustrating that adaptation involved both the persistence of some adaptive mutations and the replacement of others as selective conditions changed. Most notably, populations that initially evolved under predator exposure frequently followed different evolutionary trajectories than populations that initially evolved under predator-free conditions, despite subsequently experiencing identical alternating environments. These observations provide strong evidence that early selective history constrained subsequent adaptation by influencing which evolutionary pathways remained accessible, consistent with experimental and theoretical studies demonstrating that historical contingency can profoundly shape long-term evolution (27-29). Together, these findings indicate that fluctuating selection does more than simply alter the rate of adaptation. Instead, temporal variation in predator pressure changes which mutations are favored, how long they remain beneficial, and which future evolutionary trajectories become accessible. Consequently, ecological context and evolutionary history interact to shape adaptive outcomes, emphasizing that the predictability of bacterial evolution depends not only on the selective environment itself but also on the sequence in which selective pressures are encountered.

Although our analyses focused on high-frequency mutations enriched for signatures of positive selection, we cannot exclude the possibility that some mutations rose in frequency through genetic hitchhiking with linked beneficial alleles rather than through direct fitness effects. Likewise, the serial transfer regime inevitably imposed periodic population bottlenecks that may have influenced the fixation or loss of individual mutations through genetic drift. However, the extensive parallel evolution observed across independent populations, together with the consistent differences between constant and fluctuating predator treatments, indicates that these stochastic processes are unlikely to account for the major evolutionary patterns reported here.

Hypermutators arise through defects in DNA mismatch repair genes, most commonly *mutS* or *mutL*, resulting in elevated mutation rates that increase the supply of genetic variation available for selection (8,30). Hypermutator strains are frequently recovered from chronic *P. aeruginosa* infections, where they facilitate adaptation to antibiotics, host immunity, and other persistent selective pressures (8,31,32). The repeated, independent emergence of hypermutators in our experimental populations demonstrates that this adaptive strategy is not restricted to clinical settings or host-associated selection. Instead, elevated mutation rates appear to represent a broadly employed evolutionary strategy that enables *P. aeruginosa* to adapt to diverse selective pressures. Hypermutator lineages have likewise evolved repeatedly in the LTEE, where defects in DNA repair genes accelerated adaptation by increasing the supply of beneficial mutations while simultaneously promoting the accumulation of numerous hitchhiking and deleterious mutations (33-36).

Despite accumulating hundreds of mutations, the three hypermutator lineages did not exhibit increased virulence in the invertebrate host model. This finding is consistent with observations from clinical populations, where hypermutator strains often display reduced rather than enhanced virulence despite their elevated mutation rates (37-39). Rather than producing increasingly pathogenic lineages, hypermutation appears to facilitate rapid adaptation to local selective pressures while simultaneously generating trade-offs that can diminish virulence. Our findings support the view that hypermutation primarily promotes evolutionary flexibility (40-42), with its phenotypic consequences depending on the selective pressures encountered during adaptation.

Beyond the emergence of hypermutators, adaptation across all treatments exhibited remarkable genetic convergence. Adaptive mutations repeatedly targeted regulatory genes involved in environmental sensing, signal transduction, motility, biofilm formation, and stress responses, including *phoQ, gacS, fleQ, morA, marR, mexT*, and *vfr*. Many of these genes are also recurrent targets during chronic *P. aeruginosa* infections and experimental evolution studies (13,43-46). Because these regulators control large downstream transcriptional networks, relatively few mutations may generate widespread phenotypic consequences, making them efficient evolutionary targets in complex and changing environments. Similar patterns of genetic convergence have emerged from the LTEE, where independent *E. coli* populations repeatedly acquired mutations in many of the same genes and functional pathways despite evolving independently for tens of thousands of generations (19,34,36). The repeated evolution of these loci across environmental, laboratory, and clinical settings suggests that adaptation in *P. aeruginosa* is constrained to a limited set of highly accessible regulatory pathways despite substantial ecological differences among habitats.

Although adaptive evolution repeatedly targeted many genes previously associated with virulence regulation, the resulting phenotypic outcomes were highly variable among evolved lineages and did not consistently align with evolutionary treatment. Despite evidence of parallel evolution at the genomic level, individual isolates exhibited diverse changes in motility, biofilm formation, siderophore production, cell morphology, and competitive performance. This phenotypic diversity contrasts with the repeated targeting of common adaptive pathways, suggesting that similar genetic targets can give rise to multiple phenotypic outcomes through different combinations of mutations and genetic backgrounds. Together, these findings demonstrate that ecological context shapes not only which mutations are selected but also how those mutations are translated into organismal phenotypes.

Perhaps the most important implication of these results is that extensive genomic and phenotypic diversification did not produce a predictable increase in virulence. Across both non-hypermutator and hypermutator populations, predator-mediated selection generated widespread changes in traits commonly associated with pathogenicity, including motility, biofilm formation, siderophore production, hemolysis, growth, and competitive ability. Yet these changes consistently failed to increase virulence in the invertebrate host model. Although these results were consistent in our invertebrate infection model, the consequences of predator-mediated adaptation for virulence in mammalian hosts remain to be determined. Nonetheless, rather than supporting a simple interpretation of the coincidental evolution hypothesis in which environmental predation directly promotes pathogenicity, our findings suggest that predator-mediated selection primarily restructures the adaptive landscape from which pathogenic phenotypes may eventually emerge. Virulence therefore appears to arise through complex interactions among environmental adaptation, genetic background, and host-specific selective pressures rather than representing an inevitable consequence of adaptation to environmental predators (14,16,47).

## Methods

### Experimental evolution

Experimental evolution was initiated as described previously (15). Briefly, six replicate populations of *P. aeruginosa* NRB were propagated in Neff medium either alone (media-evolved; ME) or in the presence of *Tetrahymena thermophila* (predator-evolved; PE) by daily 4% transfers at 35°C. At Day 60, each of the six replicate ME and PE populations was split into two parallel populations. One set of populations continued under the original conditions (ME or PE), whereas the second set founded the fluctuating treatments by transferring descendants of the ME populations to predator-containing medium (MP) and descendants of the PE populations to predator-free medium (PM). MP and PM populations alternated between predator-present and predator-free environments every two weeks until Day 180 (∼828 generations), whereas ME and PE populations continued under their original conditions.

Predator viability was monitored daily by light microscopy, while bacterial persistence was confirmed by plating on Luria–Bertani (LB) agar. Both species remained detectable throughout the experiment. At Days 120 and 180, populations were plated on LB agar, and one colony was randomly selected from each replicate population for whole-genome sequencing and downstream phenotypic analyses. Isolates were stored at −80°C in 20% glycerol.

### DNA extraction, sequencing, and genome analysis

Genomic DNA was extracted from overnight cultures using the Zymo Quick-DNA™ Miniprep Plus Kit (Zymo Research) according to the manufacturer’s instructions. Sequencing libraries were prepared using the Illumina Nextera DNA Flex Library Preparation Kit and sequenced on an Illumina MiSeq100 platform (2 × 150 bp paired-end reads). Both evolved populations and individual isolates were sequenced.

Sequencing reads were quality trimmed using Trimmomatic v0.40 (48) and mapped to the ancestral genome using breseq v0.39.0 (49). Only bases with a PHRED quality score ≥30 were considered, and variants were called using a minimum allele frequency threshold of 5%. Average sequencing coverage and mapping statistics are provided in **Table S1**. Variants identified when ancestral reads were remapped to the reference assembly were considered sequencing or assembly artifacts and excluded from all downstream analyses.

Neutral expectations for mutation classes and parallel evolution were estimated using a custom Python simulation. Briefly, the mutagenicity and mutation spectrum of each nucleotide was estimated from the observed variants across all lines. Using these estimates, we conducted 10,000 independent simulations where the observed number of mutations for each line was randomly introduced into the ancestral genome while preserving the relative mutability and mutation spectrum of each nucleotide. Genome annotations were used to classify mutations as intergenic, synonymous, or nonsynonymous, and to estimate the expected frequency of gene- and site-specific parallel evolution under neutrality.

### Selection coefficient estimation

Selection coefficients ((s)) were estimated from changes in individual allele frequencies across consecutive 60-day evolutionary intervals (Days 0–60, 60– 120, and 120–180). Under our experimental conditions, each interval corresponded to approximately 276 bacterial generations. Selection coefficients were calculated from logit-transformed allele frequencies as:

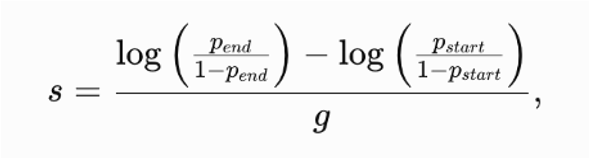

where *p*_start_ and *p*_end_ represent allele frequencies at the beginning and end of each interval, respectively, and *g* is the number of generations (276). Positive values indicate increasing allele frequencies, whereas negative values indicate declining frequencies (**Dataset S2**).

Only mutations reaching an allele frequency ≥5% in at least one sampled population were retained. Because the logit transformation is undefined for allele frequencies of 0 and 1, mutations below the 5% detection threshold were conservatively assigned a frequency of 0.05, whereas fixed mutations were assigned a maximum frequency of 0.99 prior to calculating selection coefficients. Hypermutator populations were excluded from this analysis.

Selection coefficients were calculated independently for each mutation within each replicate population. For fluctuating treatments (MP and PM), the Day 0–60 interval used the corresponding ME or PE populations because these populations shared identical evolutionary histories before environmental switching. Subsequent intervals were calculated using allele frequencies measured directly from MP and PM populations.

Because selection coefficients were estimated across 60-day intervals, they represent interval-averaged estimates of selection. Consequently, mutations that rapidly swept to fixation within an interval may have experienced stronger instantaneous selection than reflected by the estimated values. Furthermore, these estimates describe the net change in frequency of the lineage carrying each mutation and should not be interpreted as direct estimates of the fitness effects of individual mutations, as linked selection and clonal interference may also contribute to observed allele-frequency dynamics.

### Phenotypic assays

Unless otherwise stated, overnight cultures of the ancestral strain and evolved isolates were grown in Neff medium, standardized to OD_600_=1, and used for all phenotypic assays. One isolate (PM2 Day 180) was excluded from phenotypic analyses because of suspected contamination. In addition, the three hypermutator isolates were analyzed separately from the non-hypermutator isolates. Growth in Neff medium, growth in the presence of *Tetrahymena thermophila*, growth in germ-free honey bees, swimming motility, and swarming motility assays were performed as previously described (15) using three independent biological replicates. Biofilm formation, extracellular protease activity, siderophore production, and hemolytic activity assays were also performed as previously described (15) using two independent experiments with four technical replicates per experiment. Virulence was assessed in germ-free honey bees as described previously (15) by feeding newly emerged bees bacterial suspensions in sterile sugar syrup and monitoring mortality every 24 h for five days. Virulence assays were performed in two independent experiments with three biological replicates per experiment.

### Competition assays

Competition experiments were conducted under three environments: Neff medium alone, Neff medium containing *T. thermophila*, and germ-free honey bees. For each assay, overnight cultures of evolved isolates and the ancestral strain were standardized to OD_600_ = 1 and mixed at equal starting abundances. Competition mixtures included pairwise competitions between each evolved isolate and the ancestor as well as mixed competitions among evolved populations and the ancestor.

For in vitro competitions, mixtures were inoculated into either Neff medium or predator-containing cultures and incubated for 24 h. For in vivo competitions, newly emerged germ-free honey bees were fed standardized bacterial mixtures and maintained under hive-mimicking conditions for 20 h before gut dissection.

Following each competition assay, bacteria were recovered by plating on cetrimide agar. Colonies from countable plates were pooled, genomic DNA was extracted, and whole-genome sequencing libraries were prepared as described above. Relative abundances of competing isolates were determined from isolate-specific genomic variants identified by whole-population sequencing.

### Scanning electron microscopy

The ancestral strain and all Day 180 isolates were grown for 48 h in Neff medium before fixation in 2% paraformaldehyde and 2.5% glutaraldehyde. Cells were mounted on poly-D-lysine-coated coverslips, dehydrated through an ethanol series, critical-point dried, sputter-coated with gold/palladium, and imaged using a Zeiss Supra 25 field-emission scanning electron microscope at the Microscopy Services Laboratory, Department of Pathology and Laboratory Medicine, UNC Lineberger Comprehensive Cancer Center. (https://www.ebi.ac.uk/biostudies/bioimages/studies/S-BIAD2916).

Cell area was quantified from SEM images using ImageJ. Images were processed using median filtering, thresholding, and particle analysis. Between 97 and 137 cells were measured per isolate, after which 95 cells were randomly selected from each isolate using the Python random.sample() function to standardize sample size across treatments prior to statistical analyses.

## Acknowledgements

We would like to thank Kristen K White, Jillann A Madren and Victoria J Madden from The Microscopy Services Laboratory, Department of Pathology and Laboratory Medicine, supported in part by P30 CA016086 Cancer Center Core Support Grant to the UNC Lineberger Comprehensive Cancer Center. We would also like to thank Dr. Nikhil Bardeskar for his assistance with the virulence assays. We are grateful to Dr. Parul Bardeskar for her assistance with bee and SEM assays. Lastly, we thank Dr. Louis-Marie Bobay for his feedback and editorial suggestions of the manuscript.

This work was supported by the Research Capacity Fund (HATCH), project award no. 7006801, from the U.S. Department of Agriculture’s National Institute of Food (to K.R.) and Agriculture and the National Institutes of Health under grant 7R01GM145747 (to K.R.).

