## Supplemental Figures and Tables for "Parallel and Divergent Evolution in *Pseudomonas aeruginosa* Under Constant and Fluctuating Predator-Mediated Selection"

**Supplementary Material**

*Supplementary Figures*

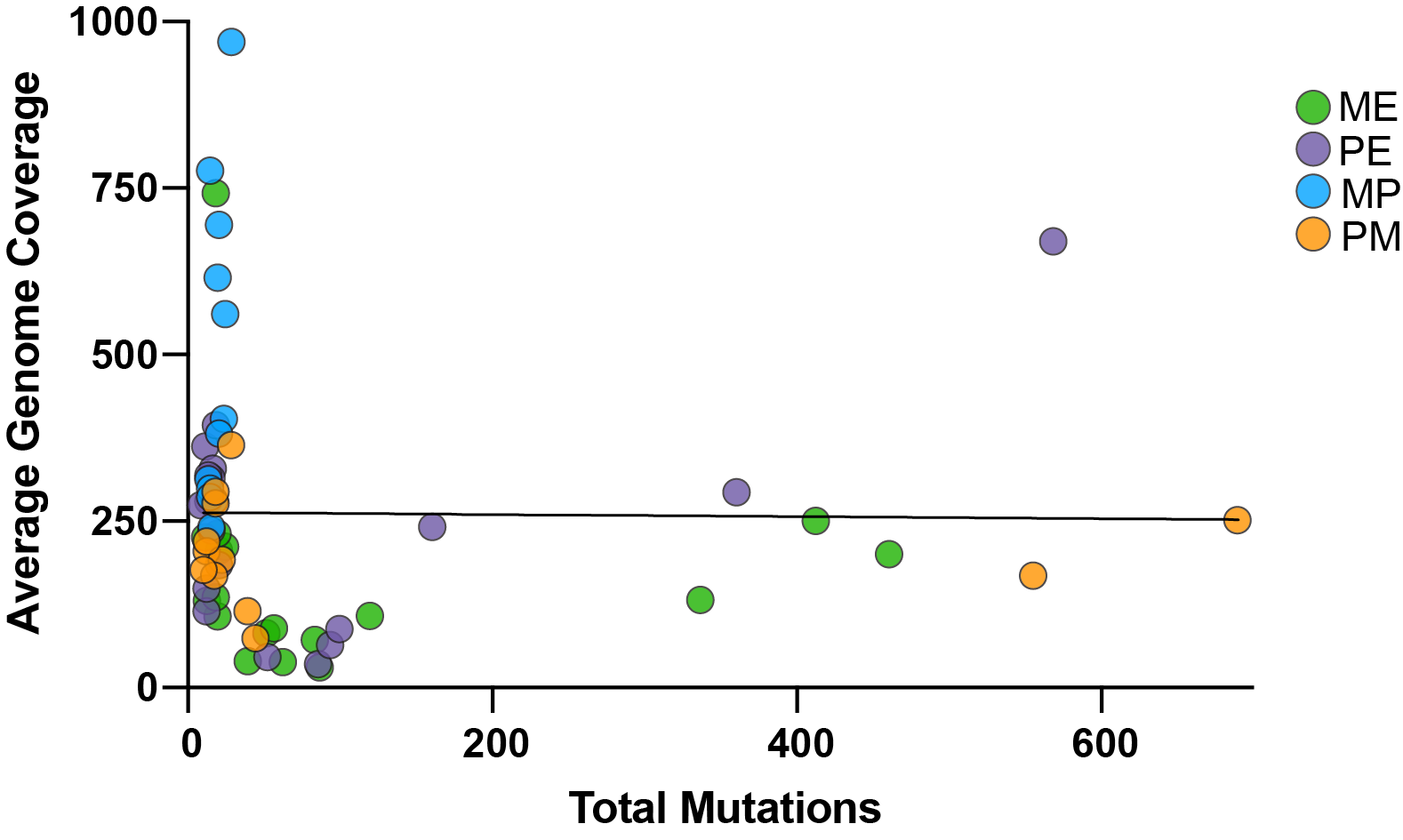

**Figure S1.** Relationship between the total number of mutations per population at Days 60, 120, and 180 and average genome coverage. *R^2^=*0.0001498, *P=*0.9260

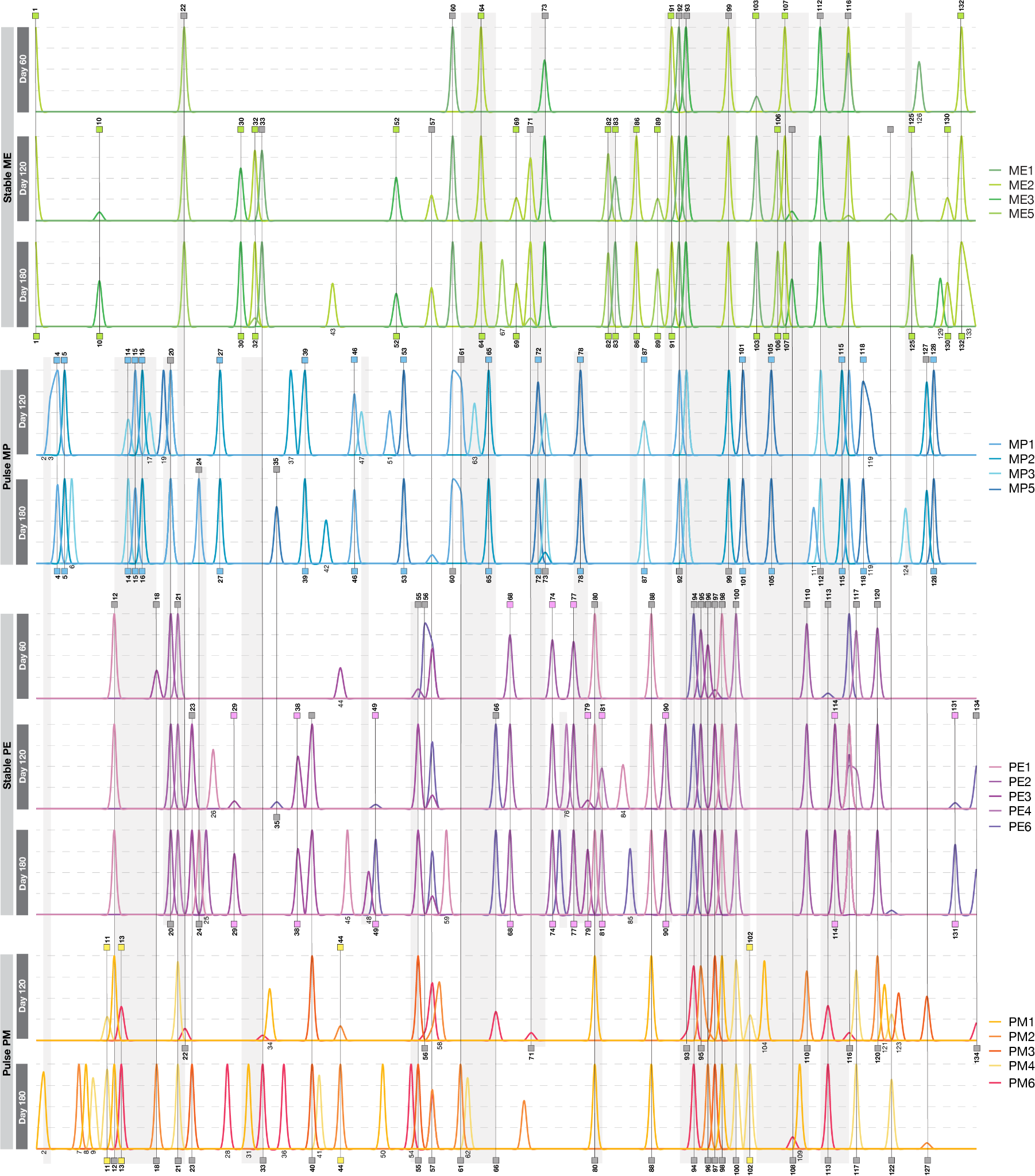

**Figure S2**. High-frequency mutations across stable and fluctuating evolutionary treatments (excluding hypermutators). The x-axis shows individual mutations reaching ≥0.5 frequency in at least 1 population, arranged by position in the genome. The y-axis represents mutation frequency (0–1). Vertical lines highlight same-site mutations. Gray shading represents same-gene mutations. Mutation numbers correspond to **Dataset S3.**

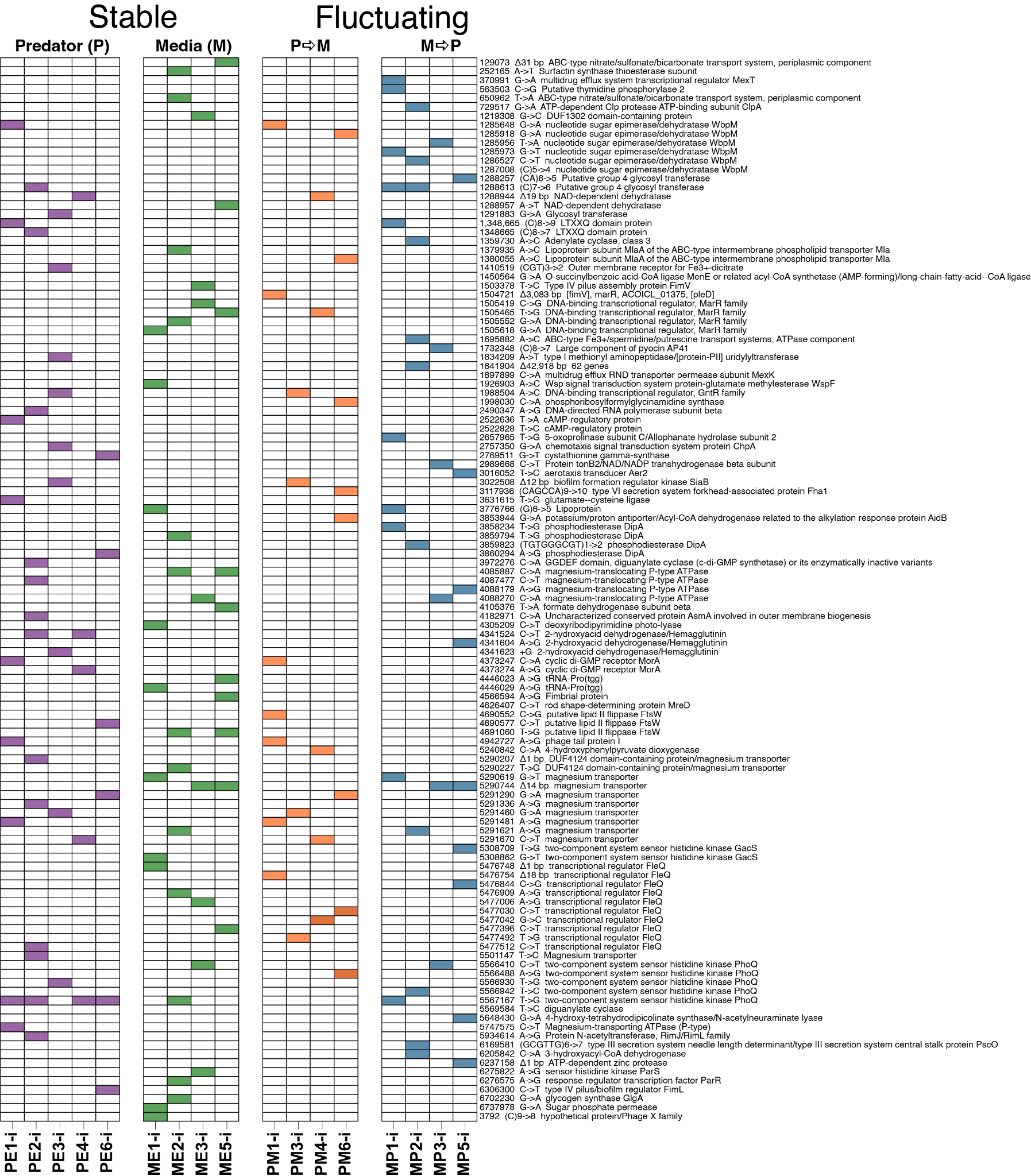

**Figure S3.** Mutations identified in the tested isolates. Detailed mutation information is provided in **Dataset S4**.

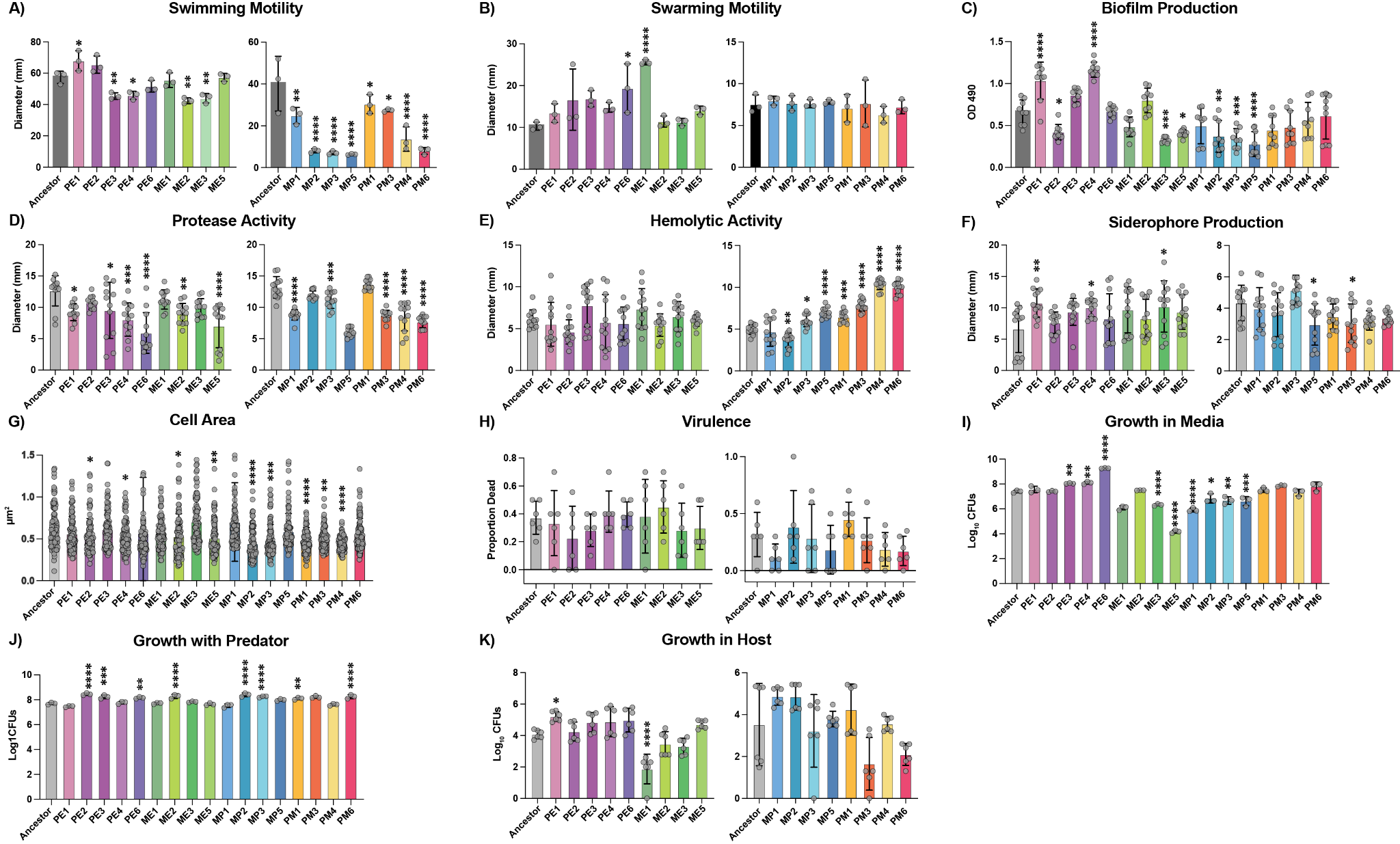

**Figure S4. Phenotypic characteristics of evolved isolates.** Replicate phenotypic assays were performed for isolates from each evolved line (hypermutators excluded) and compared to the ancestral strain for virulence-associated traits and growth. Assays included: (A) swimming motility, (B) swarming motility, (C) biofilm production, (D) protease activity, (E) hemolytic activity, (F) siderophore production, (G) cell area, (H) virulence in honey bees, (I) growth in media, (J) growth in co-culture with *T. thermophila*, and (K) growth in honey bees. Statistical significance was assessed using a Kruskal–Wallis test followed by Dunn’s multiple comparisons test. For each assay, evolved isolates were compared individually to the ancestral strain. Significant differences relative to the ancestor are indicated by asterisks (* = P <0.05, ** = P <0.01, *** = P <0.001, **** = P <0.0001). Data are summarized in **Figure 4.**

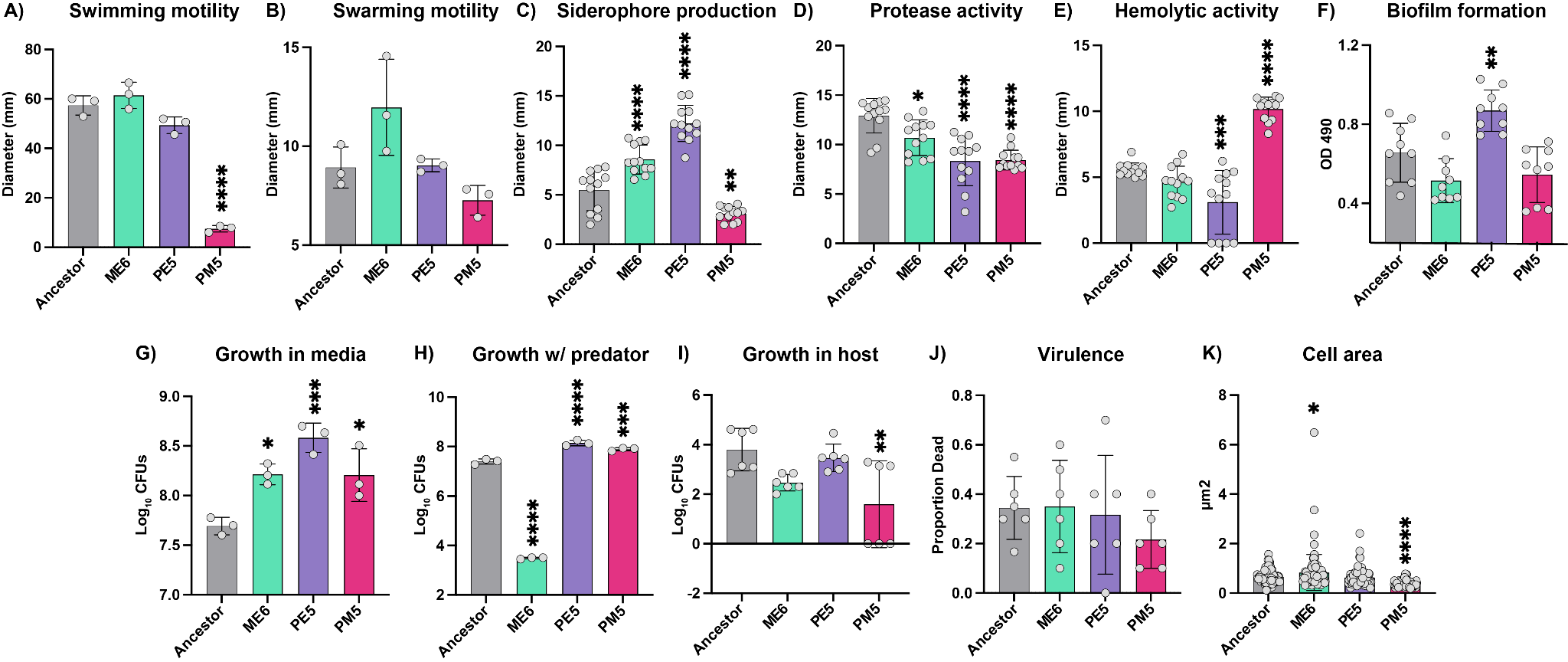

**Figure S5. Evaluation of phenotypes of hypermutator strains.** Replicate phenotypic assays were performed for isolates from each hypermutator-evolved line and compared to the ancestral strain for virulence-related phenotypes and growth traits. Assays included A) swimming motility, B) swarming motility, C) siderophore production, D) protease activity, E) hemolytic activity, F) biofilm production, G) growth in media, H) growth in co-culture with *T. thermophila* (growth w/ predator), I) growth in honey bees (growth in host), J) virulence in honey bees, and K) cell area. Statistical significance was assessed using a Kruskal–Wallis test followed by Dunn’s multiple comparisons test. For each assay, individual evolved isolates were compared to the ancestral strain. Significant differences relative to the ancestor are indicated as * (P < 0.05), ** (P < 0.01), *** (P < 0.001), and **** (P < 0.0001). Results are summarized in **Figure 6.**

*Supplementary Tables*

**Table S1:**  Average genome coverage of all isolates and populations with alignment percentages.

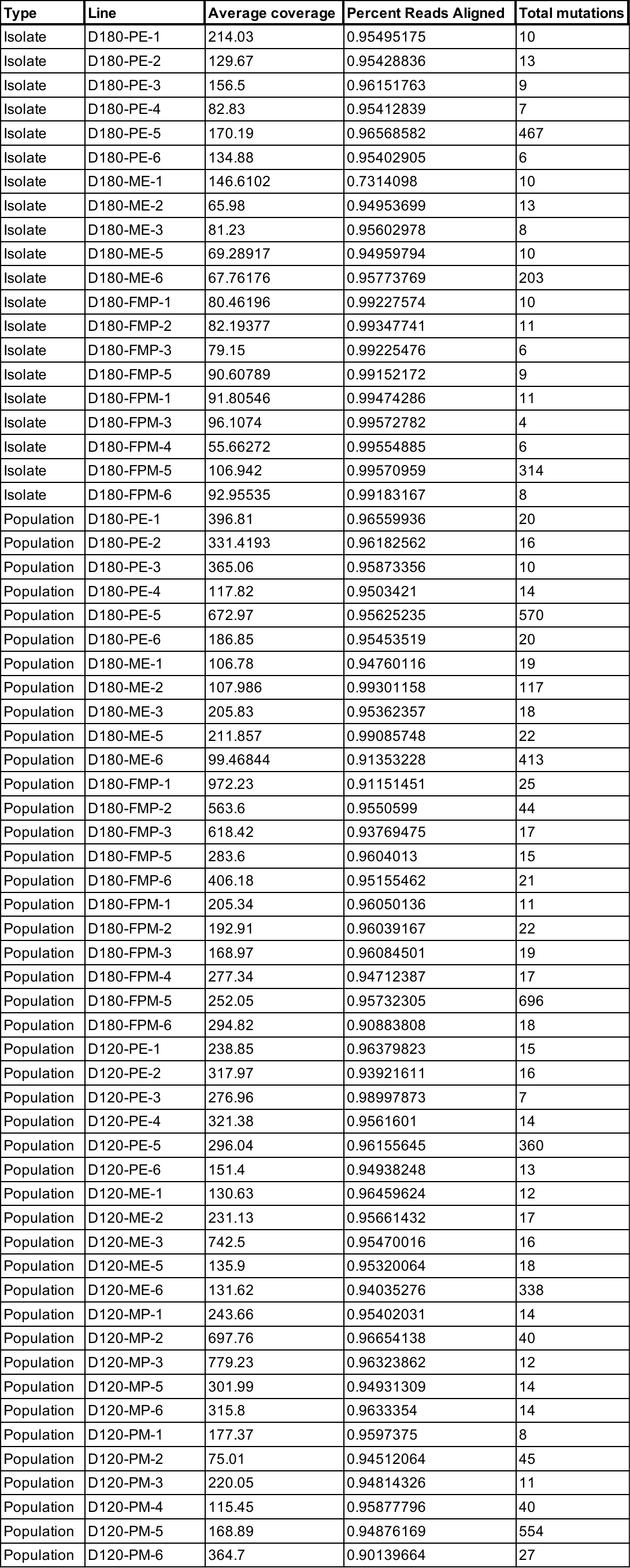

**Table S2:** Proportion of mutation types

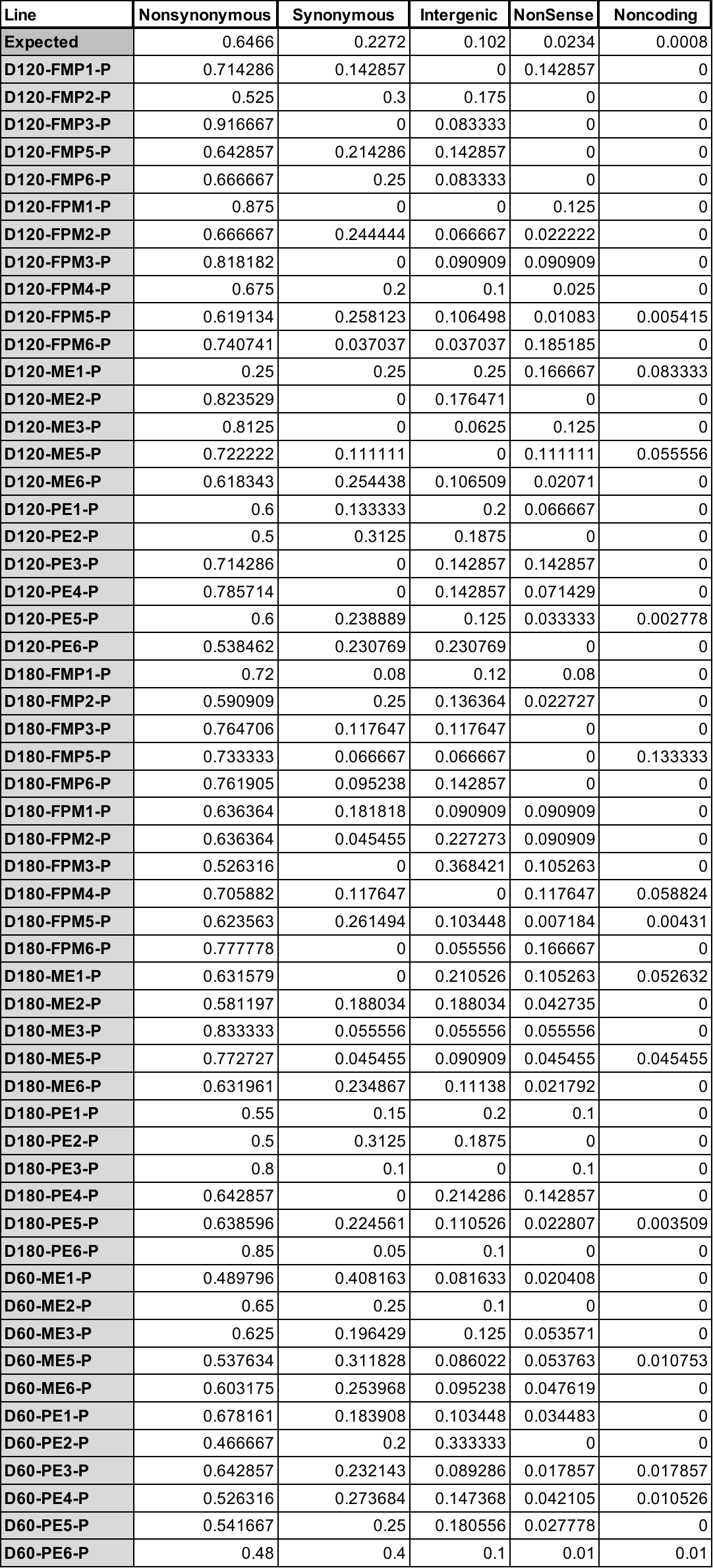

*Supplementary Datasets*

**Dataset S1**. Mutations identified in evolved populations. List of mutations detected across all evolved populations at a frequency greater than 5%.

**Dataset S2**- Selection coefficient results.

**Dataset S3.** High-frequency mutations reaching a frequency of ≥50% in evolved populations excluding hypermutator lines. The number of each mutation corresponds to the numbers in **Figure S2**.

**Dataset S4.** Mutations identified in evolved isolates (excluding hypermutators). List of mutations detected across all evolved isolates at a frequency greater than 5%.

**Dataset S5** Mutations identified in hypermutator evolved isolates.
